# Divergent climate impacts despite similar response to temperature in a widespread aerial insectivore

**DOI:** 10.64898/2025.12.15.694424

**Authors:** C. C. Taff, J. R. Shipley, D. R. Ardia, D. Aborn, L. Albert, M. Bélisle, A. Belmaker, L. L. Berzins, T. Blake, F. Bonier, H. C. Brewer, M. W. Butler, K. Cameron, S. B. Case, D. Chang van Oordt, R. G. Clark, E. D. Clotfelter, A. R. Cox, R. D. Dawson, E. P. Derryberry, A. M. Diaz Bohorquez, P. O. Dunn, V. Ferretti, A. M. Forsman, M. Fuirst, D. Garant, D. R. Garrett, J. Gutiérrez, J. C. Hagelin, B. M. Hardt, M. E. Harris, K. Horton, C. Houle, J. L. Houtz, P. L. Jones, K. C. Jordan, A. S. Kindel, R. Klaver, S. A. Knutie, K. S. Lauck, M. P. Lombardo, S. C. Lougheed, A. C. Love, S. A. Mackenzie, J. P. McCarty, A. E. McKellar, N. Mejia, C. A. Morrissey, M. L. Nahom, D. R. Norris, L. M. Para, F. Pelletier, C. K. Porter, W. B. Rendell, E. A. Riddell, J. W. Rivers, R. J. Robertson, A. Rose, K. A. Rosvall, T. A. Ryan, R. P. Shannon, D. Shutler, V. F. Simons, M. Stanback, C. E. Tarwater, P. A. Thorpe, M. W. Tingley, C. L. Tischer, B. A. Tonelli, M. L. Truan, C. W. Twining, J. J. Uehling, C. Vleck, D. Vleck, M. L. Watson, N. T. Wheelwright, L. A. Whittingham, D. W. Winkler, C. Youngflesh, C. Zimmer, M. N. Vitousek

## Abstract

Climate change is shifting when animals breed^1,2^, but it is still not clear why some populations keep pace with warming while others fall behind^3,4^. Differences could arise from variation in sensitivity to temperature^3^ or constraints on the ability to respond to temperature. Without knowing whether populations differ in sensitivity—or in their ability to act on that sensitivity—we cannot identify which are most at risk. Using 1,555 population-years from 123 populations of tree swallows (*Tachycineta bicolor*), we show that populations have similar sensitivity to local temperature, advancing breeding by about one day per degree of warming. However, northern populations face tighter time constraints and greater exposure to recent warming. Northern populations have advanced laying dates the most, but still experience stronger selection for earlier breeding, especially in warm years; they have also declined most in breeding abundance. These findings show that vulnerability to climate change can arise not just from different sensitivity to warming, but from when and where populations can respond effectively. By disentangling sensitivity from timing constraints, our results support a general mechanism by which even uniformly responsive species can show uneven impacts of climate change across their ranges.

---

Phenological shifts are among the clearest consequences of climate change^5^ and advances in the timing of annual life history events, such as breeding dates in birds, have been well documented^1,6,7^. Although earlier breeding is common, taxa vary considerably in the degree to which timing tracks—or fails to track—temperature changes^8–10^. Among species, these differences in plasticity or rapid evolution can drive variation in vulnerability to climate change^10–12^. However, for broadly distributed species, similar variation in phenological sensitivity—defined here as the slope of egg laying date to temperature variation^3^—among populations may result in geographic differences in climate vulnerability. Yet relatively little is known about geographic variation in phenological sensitivity in most taxa^3,4,13,14^.

Among vertebrates, very few species have the long-term, geographically replicated data required to study range-wide variation in phenological sensitivity of breeding dates^3,4,15^ and species-level estimates of sensitivity often rely on data from a single or few populations per species^11^. Relying on a single location is problematic because it cannot capture spatial variability in phenological sensitivity or vulnerability. Extrapolating sensitivity estimates from one population could overestimate vulnerability by, for example, failing to account for local adaptation in the way cues are used to time breeding^16^. Alternatively, erosion of variation in phenological sensitivity in populations most exposed to climate change could lead to an underestimation of range-wide vulnerability^3^. Finally, the same behavioral response to temperature could still result in divergent climate impacts when populations differ in climate exposure, morphology, or timing constraints^7,17,18^, resulting in a difference in the effectiveness of responses to temperature. Thus, understanding range-wide intra-specific patterns is crucial because variation in vulnerability could exacerbate or ameliorate the impacts of climate change and complicate efforts to mitigate it.

Here, we studied phenological sensitivity and climate impacts in tree swallows (*Tachycineta bicolor*) using 94,873 nests from 123 population time series spanning 31 degrees of latitude combined with estimates of arrival timing, population abundance trends, and historical temperature exposure over the past half century (Figure 1). We found that tree swallows across their range lay eggs earlier in warmer springs, with each 1°C increase in temperature leading to a nearly one-day advancement in egg laying (Figure 2A; Table S1; β = −0.95, CI = −1.05 to −0.85 days, P < 0.001). The temperature period that most strongly influences breeding timing (i.e., the sensitive window^3^) occurs during the three weeks before females typically lay, suggesting that cues experienced shortly before breeding drive phenological adjustments (Figure S1; the best supported temperature window opened 21.1 ± 3.7 days, mean ± SD, before population-specific mean laying date and closed 0.1 ± 3.6 days after mean laying date). Phenological sensitivity was similar across populations spanning the species’ range—from southern to northern sites and across different habitat types—indicating that tree swallows exhibit a broadly conserved thermal response in reproductive timing despite variation in local thermal regimes (Table S1-S2).

**Figure 1.**
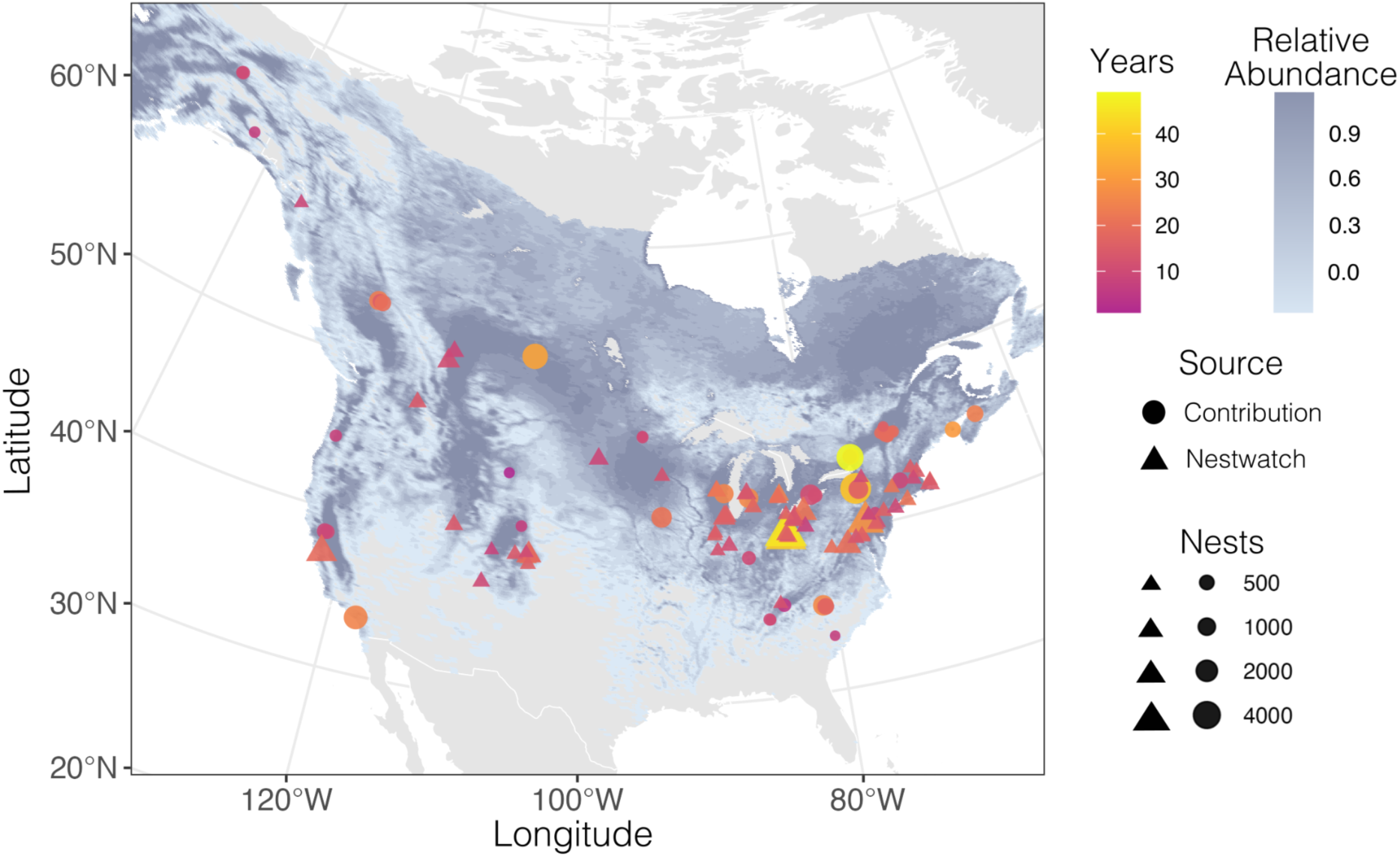
Spatial and temporal distribution of sampling from 123 populations across 31 degrees of latitude. Color gradient indicates the number of years of observation at each site. Size and shape of each point indicate the total number of nests and whether the data were contributed by a research group or extracted from NestWatch. The range map shows estimated relative abundance during the breeding season from eBird status and trends, which denotes relative abundance as the expected number of birds observed on a standard eBird checklist. Background map is from rnaturalearth and the plot is produced in Albers equal area projection.

**Figure 2.**
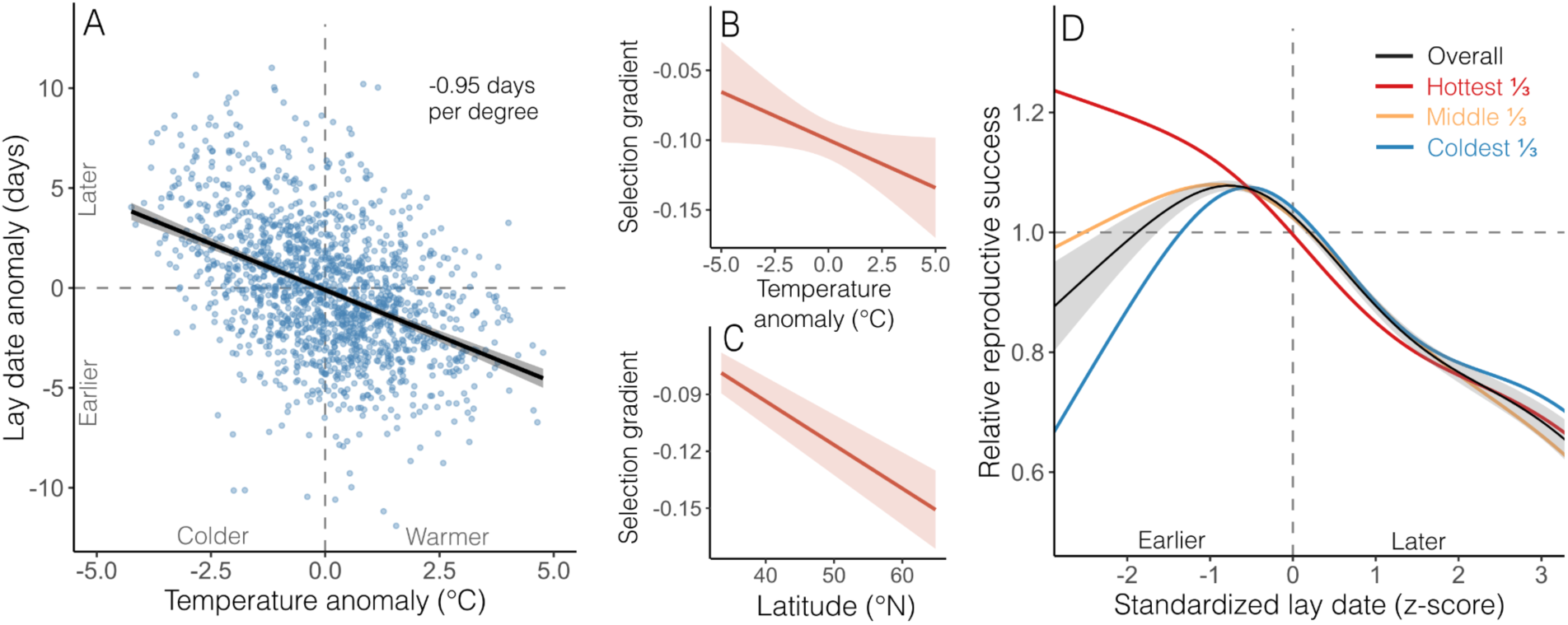
Phenological sensitivity to temperature and selection on timing of breeding. (A) Overall relationship between population specific temperature anomaly and laying date anomaly (anomalies are calculated as yearly deviations from long-term population averages). Each point is one population-year, and the fit line is from a mixed model accounting for spatial autocorrelation and population with CI in gray. (B-C) Relationship between the within-population-year standardized linear selection gradient on timing of breeding and yearly temperature anomaly (B) or latitude (C). Patterns shown are derived from a GAMM accounting for spatial autocorrelation and with all parameters held at their mean values. (D) Relationship between standardized lay date and relative reproductive success across all populations accounting for spatial autocorrelation. Black line and shaded region show the overall fit and CI while red, orange, and blue lines show the fit when splitting to the highest, middle, and lowest thirds of the relative temperature anomaly distribution.

Similar phenological sensitivity across the range contrasts with the most comparable studies of among population variation in other species; studies of European cavity nesting species demonstrated population specific variation in phenological sensitivity in blue (*Cyanistes caeruleus*) and great (*Parus major*) tits and collared (*Ficedula albicollis*) and pied (*Ficedula hypoleuca*) flycatchers^3,4^. Overall, local residents (tits) responded more strongly to temperature than long distance migrants (flycatchers) and it has been suggested that the timing constraints imposed by migration drive these differences^4^. Tree swallows have an intermediate migration strategy that, in some parts of the range, involves relatively short migration with a long pre-breeding stage^19^. Despite this life history, we still found that tree swallows had weaker mean phenological sensitivity (−0.95 days per °C) than reported in all four European species (days per °C: blue tit = −3.4, great tit = −2.8, collared flycatcher = −1.5, pied flycatcher −1.5)^3,4^. Studies that have compared the phenological sensitivity of many species at single locations have found a wider range of responses, but could not test for among population variation^11^. Our results highlight the need for studies on a wider range of species with different life histories to understand both intra- and inter-specific variation in climate change vulnerability.

Despite adjusting egg laying dates in warmer springs, populations still experienced consistent selection for breeding even earlier (Figure S2; standardized linear selection gradient from 85 populations and 832 population-years with at least 30 monitored nests = −0.10; CI = −0.11 to −0.08). Selection was apparent across the range and did not differ with habitat type (land use PC1: *t* = −0.03, P = 0.97; land use PC2: *t* = −0.92, P = 0.36). Selection for earlier breeding is well documented in many birds^11,20^, including tree swallows^21^. Ongoing selection for early breeding is consistent with an inability for rapid evolution or plasticity to keep pace with climate change^11^, but apparent selection can also persist at equilibrium when non-heritable differences, such as nutritional state, influence both the timing of breeding and reproductive output^20^. However, we also found that the linear selection gradient for early breeding was stronger in northern populations and in relatively warmer years (Figure 2B-C; Table S3; latitude *t* = −2.02, P = 0.04; temperature anomaly *t* = −14.03, P < 0.001). Thus, our results indicate that at least some of the selection for earlier breeding was mediated by climate exposure.

Pooling nests from all populations in a single model revealed an overall pattern of stabilizing selection (Figure 2D; Table S4; smoothed term for standardized lay date F = 338.0, P < 0.001), but the qualitative pattern of selection depended on relative temperature during the sensitive window. In years that were cold relative to population averages, we observed a clear pattern of stabilizing selection on laying date, but in warm years we observed directional selection for earlier laying (Figure 2D). Overall, the maximum relative reproductive success was achieved in nests that were initiated 2.7 ± 1.2 days before the population mean laying date. Our findings demonstrate that variation in early season temperature was an important mediator of reproductive success. Across all the sites and years that we studied, year-to-year temperature variability during the sensitive window was higher in northern populations (GAM with a linear effect of latitude β = 0.13 ± 0.015, t = 9.1, P < 0.001). As a consequence, earlier breeding—especially in northern populations—can increase the probability that breeders are exposed to cold, wind, and precipitation, resulting in nest failure in some years^22–24^. In contrast, recent work suggests that tree swallows handle heat exposure relatively well^25^. Thus, timing constraints associated with migration combined with variation in local cues may be especially critical for this species^26^.

Conflicts between migration and breeding timing could constrain adaptation to climate change when climate change exposure differs on the non-breeding grounds, migratory route, and breeding grounds^7^. The severity of this constraint could vary geographically because northern populations typically have longer migration distances and further advances in egg laying could be limited by migratory arrival timing. Tree swallow migration is highly sensitive to spring green up and arrival timing varies considerably among years^17^. However, when integrating arrival dates with our monitored populations, we found that relatively early migration arrival did not result in relatively early egg laying, as these dates were uncorrelated within populations (GAMM with random effect for population and controlling for spatial autocorrelation; *n* = 1,212 site-years, linear effect of arrival anomaly on laying anomaly β ≥ −0.1, CI = −0.05 to 0.04, *t* = −0.2, P = 0.85). Citizen science projects have been invaluable in detecting the macro-ecological effects of climate change, such as large-scale movements^10,17,27^; our result highlights the value in linking these datasets with monitoring records of individual animals to connect variation in breeding timing with individual level consequences for reproduction.

Local decoupling of arrival timing and egg laying suggests that even species that can adjust migration timing may face constraints on optimal breeding time due to later arrival. Indeed, we found that timing of all phenological events (arrival, sensitive window opening, and laying) was later at higher latitudes (Figure 3A; Table S5; P for all latitudinal smooths < 0.001), but latitudinal changes in timing of these events were not parallel. While arrival dates continued to get later at higher latitudes, both sensitive window opening dates and average egg laying dates plateaued above approximately 45°N. As a consequence, the number of days between population-specific arrival and mean egg laying date decreased at higher latitudes (Table S5; Figure 3B; P < 0.001) suggesting that the northernmost populations face a time constraint such that the scope for further advancing breeding date without missing part of the local sensitive window is limited by migration arrival timing. Comparative studies suggest that a similar constriction of pre-laying time on the breeding grounds at higher latitudes may be a common challenge for widespread species^17^.

**Figure 3.**
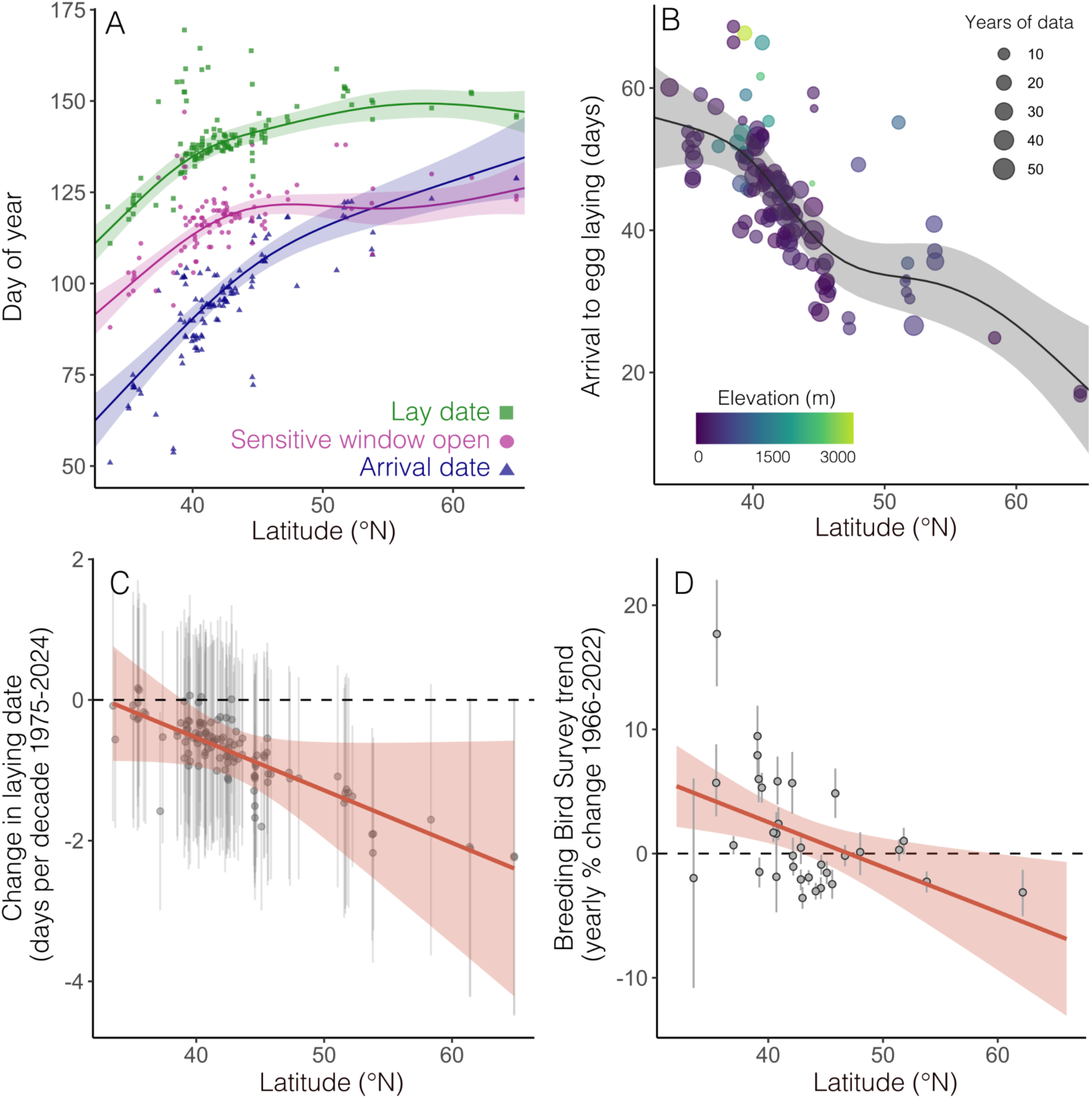
Latitudinal variation in timing of breeding season life history events, rate of phenological advancement, and long-term population trajectories. (A) Relationship between latitude and arrival date, sensitive window opening date, and laying date. Lines and CIs are from GAMMs using all years of observation, but points illustrate the overall average across all years for each site. Most outlier points are from sites > 900m a.s.l.; including or excluding these sites or adding elevation as a predictor to models does not change the qualitative pattern. (B) Relationship between latitude and the number of days between arrival and mean egg laying date. Each point is from one population with elevation and number of years indicated by color and size. The curve shown is from the GAMM described in the text that uses 1,212 population-year values. (C) Relationship between latitude and rate of change in average laying date from a GAMM controlling for spatial autocorrelation. The red line and shaded region are the overall trend and CI for the trend. The gray points and error bars are the random slope estimates and CI for each included population. (D) Relationship between latitude and state or province level estimates for the yearly percent change in breeding season abundance from the published USGS model of Breeding Bird Survey data. Red line and shading show estimated slope and CI from a simple linear model, and gray points and error bars show the state/province level estimates and CI.

Escalation of timing constraints in the sequence of life history stages at higher latitudes indicates that these populations may be facing a greater challenge from climate change. Interestingly, whereas most species have shifted their ranges poleward in response to climate change^1,5,28^, the tree swallow range has expanded southward over the past few decades^29,30^ and some well-monitored northern populations have experienced steep declines^31^. Three lines of evidence are consistent with a latitudinal gradient in climate impacts across the populations we studied. First, we found that average temperature during our identified sensitive window dates for each population has increased universally from a reference period (1950-1974) compared to the most recent records (2000-2024) by 0.23 ± 0.13 °C; yet the magnitude of this warming was greater at higher latitudes (Table S6; latitude *t* = 35.3, P < 0.001). Second, across all populations we monitored, average date of egg laying has changed by −0.82 ± 0.12 days per decade since 1975, but the rate of advancement has been faster at higher latitudes (Figure 3C; Table S6; interaction between year and latitude *t* = −3.0, P = 0.003). As a consequence, we see little change in laying dates over the past half-century at the southernmost populations, but up to a 10-day advance in the northernmost populations. Finally, linking the location of our monitored populations to state and province level estimates of breeding season abundance over a similar time range (1966-2022)^32^ revealed a latitudinal gradient, with populations in the south generally increasing and northern populations decreasing (Figure 3D; Table S6; latitude *t* = −2.7, P = 0.01)^29^.

Taken together, we found that northern populations have been most exposed to a rapidly changing climate during a critical period of the full annual cycle. Despite making the largest adjustments to breeding dates, northern populations have also declined the most. While aerial insectivore declines have been linked to a variety of factors^33–36^, our results are consistent with the idea that timing conflicts impose an additional constraint on northern tree swallow populations that is exacerbated by climate change. Due to this timing constraint, northern populations have less time to recover from migration, secure a breeding site, and gain food resources before the start of laying. Moreover, the limited scope for further adjustment suggests that many populations may be approaching tipping points due to a lack of remaining flexibility to adjust timing. Climate change exposure increases with latitude^37^ and phenological responses among taxa are correspondingly more extreme at higher latitudes^38^. Yet most species ranges are shifting to higher latitudes despite this increased exposure^1,5,28^.

Relative to other species, tree swallows may be particularly vulnerable to timing constraints because of their specific life history or ecology. Across most of their breeding range, tree swallows produce a single nesting attempt per season and do not renest if an early nest fails^39^. As obligate aerial insectivores, tree swallows are also entirely dependent on local flying insect availability, which can shift rapidly with local temperature conditions^40,41^. We primarily focused on mean temperature, but we also found that northern populations experienced more year-to-year variation in temperature and future changes to temperature variability and precipitation might produce an additive effect to the pattern we found^22,23,42^. While mean temperature alone can have a causal effect on breeding timing^43^, there is increasing recognition that temperature variability might be an important driver of breeding decisions and population dynamics^24,44–47^ and that the frequency and intensity of extreme climate events is increasing^48^. For tree swallows, inclement conditions can drastically reduce food availability and just a few consecutive days of these conditions can result in mass mortalities^22,23,49^. Thus, northern populations must contend with the dual challenge of both warmer springs and increased exposure to temperature variability^50^.

Overall, our results demonstrate a remarkably uniform role for temperature influencing breeding timing across a continental range. Despite similar sensitivity, constraints imposed on timing vary considerably with latitude because phenological shifts in arrival, sensitive windows, and egg laying are uneven. Thus, northern populations have less opportunity to respond effectively. Northern populations have advanced their breeding dates the most yet have still declined most in abundance over the past half century. These findings demonstrate the power of combining replicated ecological monitoring with citizen science data to reveal large-scale patterns that would not be apparent in single location or short-term studies. Even when populations respond similarly to temperature, geographic variation in climate vulnerability and resilience may be common due to differences in the opportunity to respond, but the outcome of this variation will depend on species-and population-specific life history sequences. Thus, predicting the consequences of climate change will require an understanding of inter- and intra-specific variation in both sensitivity and in the opportunity to adjust to changing climates.

## MATERIALS & METHODS

### Data contributed from monitored populations

Because of their large geographic range and the accessibility of nest boxes, tree swallows are among the most studied wild bird species in the world^51^, which makes them especially powerful for detecting large scale patterns. We solicited data contributions from researchers monitoring tree swallow populations across the United States and Canada, which encompass the entire breeding range. To identify groups, we used a combination of contacting authors from previous multi-population collaborations^52,53^, searching for more recent publications on tree swallows, and asking all contributors that we identified to forward the request to any other groups that might be monitoring tree swallows.

For inclusion in this study, we asked that groups have at least eight years of monitoring data from the same location (not necessarily in successive years) with GPS coordinates available either for each group of monitored boxes or for the exact location of each individual box. In each of those years, we required that the monitoring effort was sufficient for groups to be confident that nearly all initial nesting attempts in the nest box grid were identified with either precise first egg dates or with the ability to estimate first egg dates based on precise hatching date. We also required that no experimental treatments were applied that could have had an impact on egg laying date. For analyses that examined fledging success, we excluded nests that had manipulative experimental treatments applied after clutch initiation, but these nests were included in analyses of egg laying date. We incorporated data from some populations that had fewer than eight years of monitoring, but these data were not included in the primary analyses of phenological sensitivity.

Our request resulted in data contributions from 28 different research groups totaling 53,735 nest observations from 541 population-years of monitoring effort between 1966 and 2024. While some groups monitored sites at a single location, others monitored sites over a wider area. We split nests into distinct subpopulations by creating clusters of nest boxes within a 5-km diameter circle of each other. During the breeding season, tree swallows are central place foragers and primarily forage near the nest box with occasional trips up to a few kilometers away^54,55^. Thus, this distance represents a biologically meaningful area over which local factors could influence tree swallows during the breeding season while still allowing us to group together enough nests to accurately estimate mean laying date each year. Overall, we analyzed 60 distinct locations, resulting in a total of 738 subpopulation-years of monitoring effort (Figure 1).

### Data contributed from citizen scientists

We combined data from the contributed populations described above with nest records from NestWatch, a project managed by the Cornell Lab of Ornithology that provides publicly accessible historic nest records^56^. While NestWatch records typically do not include the same level of detailed monitoring, adding these records allowed us to fill geographic gaps from our contributed populations. NestWatch records included 107,295 nests with laying date information, but many of these observations represent idiosyncratic monitoring rather than comprehensive observation of each nest within a distinct population.

To identify clusters of NestWatch observations most comparable to our contributed populations, we used the density-based clustering of applications with noise algorithm from the ‘dbscan’ package in R^57^. We identified clusters of nests that occurred within a 5-km diameter where a minimum of 15 nest records per year were reported over a minimum of 8 years. We reasoned that these locations represented arrays of nest boxes that were monitored each year. Because some research groups also submitted their observations to NestWatch, we required that the centroid of each cluster be >15-km from the centroid of any of the contributed populations to avoid duplicate entries. This process identified 63 clusters containing 41,138 nests and 817 population-years of effort from 1977-2024 (Figure 1).

The breeding data from contributed populations and NestWatch were combined into a common format for downstream analyses. For both datasets, we attempted to filter nest records to include primarily first nesting attempts rather than late renesting or second clutches. For this filtering, we determined the date of the earliest 10th percentile of nest initiation for each population-year and then excluded any nests that were initiated >42 days after this date. While this filtering was undoubtedly imperfect, it served to exclude very late nests that often represent second attempts and have low success rates. Our final cleaned dataset included 1,555 population-years of monitoring effort from 123 sites, although the sample sizes for specific analyses described below varied after filtering.

### Daily temperature data

To estimate sensitivity of breeding timing to temperature, we downloaded daily temperature data at 2 m above ground level for each location from two different data sources: we accessed Daymet^58^ data using the ‘daymetr’ package^59^ and we accessed MERRA-2^60^ data using the ‘nasapower’ package in R^61^. Each of these sources has different pros and cons. Daymet provides high spatial resolution estimates on a 1-km grid, but hourly temperatures are not identified. Only daily minimum and maximum temperature are provided, so average temperature must be estimated assuming a symmetric temperature curve. Daymet is also not yet available for 2024 (a year in which most populations in our dataset were monitored). MERRA-2 provides average temperature estimates through 2024, but the spatial resolution is lower, with temperatures available on a 50-km grid.

We compensated for the lower spatial resolution of MERRA-2 by correcting temperatures for the difference in elevation of nest box locations and the 50-km grid average using a standard lapse rate of 6.5 °C per 1-km of elevation^5^. For this correction, we determined elevation of each site based on location using the ‘elevatr’ package in R^62^. Neither temperature product was available for the earliest nests in our dataset (Daymet and MERRA-2 begin in 1980 and 1981, respectively), but only 640 of our 94,873 included nest records occurred in 1981 or earlier, so we excluded those sites from sensitive window analyses. All conclusions were qualitatively unchanged using either climate source. We report results using MERRA-2 data because they were available for the most population-years.

For analyses of the change in temperature during the sensitive window over time and long term variability in sensitive window temperature, we used gridded daily average temperature data from the Berkeley Earth Project^63^. These data are spatially coarser (1°x1° grid), but they are available starting in 1880 and were used only to describe long term changes. We accessed the gridded global land daily temperatures from the project website (https://berkeleyearth.org/data/) and then for each population we calculated the overall average temperature during the identified 21-day long sensitive window (see below) for a baseline period (1950-1974) and a contemporary period (2000-2024). We then determined change in temperature between these two time periods for each population and used this single number to describe exposure to increased temperature during this critical window. To describe latitudinal variation in year-to-year sensitive window temperature, we calculated the standard deviation of average yearly temperature for each population location.

### Land cover data

We added land cover information to each population using the North American Land Change Monitoring System^64^. This data product classifies land cover into 19 distinct categories from Landsat satellite imagery at a 30-m resolution using consistent methods covering all of North America starting in 2005 and with updated releases every 5 years. For our analyses, we were most interested in general classification of land use type into broad categories rather than detailed land use change over time (e.g., amount of agricultural activity or amount of available open water). We chose to use the NALCMS 2010 map because this was approximately the middle of the bulk of our nest observations.

For each population in our dataset, we created a 5-km diameter circle around the centroid and used this buffer to calculate zonal statistics from the land cover type raster with the ‘exactextractr’ package in R^65^. This resulted in values for the percentage cover within the buffer for each of the 19 land use categories. To simplify these data, we combined them into 7 categories that are functionally similar with respect to tree swallow foraging (e.g., we collapsed the six distinct forest types into a single forest category). Categories we considered were: agricultural, urban, barren, forested, grass/shrub, wetland, and open water.

We further simplified land use descriptions by performing a principal component analysis (PCA) on these 7 categories and saving the first two PCs. Because the land cover data are proportional and therefore not independent of each other, we performed a center log ratio transformation on percentage values prior to PCA. After transformation, the first two principal components explained 34.1%, and 21.0% of the variance, respectively (Figure S3). Sites that scored highly on PC1 had relatively more wetland, open water, and grass/shrub whereas low PC1 scores were associated with relatively more agriculture, forest, and urban areas. Therefore, PC1 largely represented a gradient from more to less anthropogenic activity. Tree swallow nests often occur along this entire gradient, with some nest boxes along fence rows in highly agricultural areas and others in naturally open wetland and grasslands (example sites in Figure S4). Importantly, occupied sites with these characteristics often occur close together across the breeding range. In contrast, PC2 was less clearly linked to an obvious land use gradient. Sites with high PC2 scores had relatively more open water and less wetland.

### Identifying sensitive temperature windows and testing for population differences

We followed a similar approach to Bailey et al.^3^ to determine sensitive climate windows using a sliding window implemented with the ‘climwin’ package in R^66,67^. For each population, we explored a set of potential climate windows ranging from 7 weeks before until 2 weeks after the population specific mean laying date (calculated using all years of data). Climwin models were run separately for each population and used mean laying date for each year of observation as the response variable. To reduce the number of models run in each set, we only evaluated potential windows that were 2-4 weeks long and with closing dates within 2 weeks of the long-term mean egg laying date for a particular population. These restrictions did not change any qualitative conclusions compared to more inclusive settings, but they eliminated biologically implausible windows (e.g., a very short window before most birds had arrived). For each candidate window in each population, climwin calculated the average temperature across all the included days, fit a single model, and then compared all candidate models for each population by AICc^67^. The package then constructs a weighted sensitive window opening date, closing date, and slope for each population based on the relative support for each model in the set.

It is important to note that in our climwin models, each population-year is one datapoint. Thus, despite the large number of nests in our dataset, sample sizes for each sliding window analysis ranged from 8 to 44 and the sliding window approach is prone to overfitting^66^. The package can account for overfitting by implementing a simulation-based approach to determine the likelihood that an identified sensitive climate window could have been detected by chance^67^. Bailey et al.^3^ used this approach and then only retained populations for which a robust sensitive window was detected. However, this approach will necessarily exclude populations that have a smaller sample size or a shallower slope of phenological sensitivity, potentially overestimating range-wide sensitivity. Because we were interested in directly testing for population differences in sensitivity (using a random slope mixed model) we retained estimates for each population at this step.

Rather than excluding populations, we initially compared opening and closing dates of identified climate windows for all populations that had at least 8 years of data with 15 monitored nests per year. After finding consistent estimates of sensitive windows across populations and no relationship between window timing or length and latitude or land use type, we created a global ‘consensus window’ based on the average sensitive window across all populations (Figure S1). The consensus window has the advantage of a much larger dataset that reduces the influence of outlier estimates of sensitive windows from individual populations. Using the consensus window, we recalculated the average temperature experienced during sensitive periods for every population and year.

This dataset then allowed us to test for variation in phenological sensitivity across populations by fitting a single generalized additive mixed model (GAMM) with laying date as the response variable and average temperature during the consensus window as a predictor^68^. A random effect for population accounted for repeated observations and for variation in population average lay date. A random slope for the effect of temperature by population provided a formal test for variation in among population phenological sensitivity. Finally, we also included a smoothed interaction between latitude and longitude to account for potential spatial autocorrelation in sampled populations. We built on this initial model by adding land use PCs as interactions with temperature to ask whether sensitivity differed by land cover.

### Selection on laying date

For analyses of selection on laying date, we filtered the dataset to only include non-experimental nests for which number of young fledged was recorded and further limited the analysis to population-years with at least 30 observed non-experimental nests. In these analyses we considered the number of offspring successfully fledged as our measure of seasonal reproductive success. We initially calculated a standardized selection gradient for each population-year by fitting a simple model with population specific relative reproductive success (number of fledglings in a nest divided by mean success for each year and population; average = 1) as the response and within-population standardized laying date (mean = 0, sd = 1) as the predictor. These individual estimates were used to describe how frequently populations experienced selection for earlier breeding. We also used these estimates of linear selection as a response variable in GAMMs to test for associations between selection and latitude, yearly temperature anomaly during the sensitive window, and land use^69^. For those models we added the linear effects of predictors, along with smoothed effects of longitude and latitude to account for spatial autocorrelation and a random effect for population.

We next fit a global model with data from all populations to describe overall selection on breeding date across all populations and years. In this model we used population specific relative reproductive success as the response and standardized laying date centered within each population as the predictor. The much larger sample size allowed us to fit this response as a GAMM that could detect non-linear selection on laying date. We included population as a random effect along with a smooth for latitude and longitude to account for spatial autocorrelation in selection estimates. We also fit a modified version of this model stratified by yearly temperature anomaly. We split the z-score for temperature anomaly into a three-level factor to illustrate the difference in overall pattern of selection in warm, average, and cold years.

It is important to note that variation in laying date almost certainly reflects a combination of additive genetic variation and plasticity. In birds, some studies have demonstrated heritable variation in laying date^70^, while others have demonstrated either developmental plasticity influenced by early life conditions^71^, or plasticity within adulthood based on current conditions^72^. Analyses of individual tree swallow populations provide some support for the role of each of these mechanisms in generating variation in laying date^73,74^. This distinction has important implications for the predicted evolutionary response to continued climate change, but our data did not allow us to parse sources of lay date variation. Rather, our goal was simply to describe patterns of selection as an indication of a potential mismatch between population average laying dates and optimal laying dates.

### Estimated arrival dates

We incorporated estimates of arrival dates by year for the location of each population from 2002-2024 using logistic generalized additive models (GAMs) derived from eBird checklists^75^. For these analyses, population data prior to 2002 were not included because eBird data were not widely available. The modeling approach to derive population specific arrival estimates was developed by Youngflesh et al.^17^ and we closely followed the updated methods described in detail by Tonelli et al.^8^.

Briefly, the approach involved filtering eBird data to include only complete checklists of between 5 minutes to 24 hours in duration, with fewer than 11 observers, and with a distance traveled of less than 5-km. Checklists were spatially aggregated into 285-km cells and then thinned by sampling without replacement to a maximum of 5,000 records per cell/year. Filtered datasets were then used to fit logistic GAMs, from which we estimated the half-maximum date for each cell/year combination (i.e., the date on which tree swallow detection reached half of the first local maximum value). This date is less sensitive to bias than first arrival dates^17^ and represents a site-specific arrival date comparable to the site-specific mean egg laying dates derived from nest monitoring data. The approach also accounted for uncertainty in detection probability and survey effort across spatial units.

Our methods were identical to Tonelli et al.^8^, except that we included checklists from a wider date range to allow for possible detection of earlier or later arrivals (relative day of year −30 to 250) and we used a modified approach to account for sites that differed in elevation. Our data included several populations breeding at high elevations (up to 3,770 meters). To improve estimates of arrival dates for high elevation sites, we thinned checklists by elevation bands within a cell (using 500-meter elevation bands) rather than across the entire cell. This allowed us to determine half-maximum dates for each cell and elevation band for which sufficient checklists were available, which were then joined to population specific nest monitoring data. Adding arrival estimates for each site and year allowed us to examine the degree to which differences in sensitivity of egg laying date was driven by pre-laying conditions on the breeding grounds versus phenological changes in arrival timing as well as geographic differences in the length of time between arrival and egg laying.

### Time constraints between arrival and egg laying

We combined population level estimates of arrival timing (from eBird), population specific sensitive window opening dates (from sliding window analyses), and egg laying dates (from nest records) for each population to explore geographic variation in time constraints on the breeding grounds. We initially asked whether there was a linear relationship between year specific arrival date and mean egg laying date using a GAMM that accounted for population ID and for spatial autocorrelation (as described above). Next, we fit similar GAMMs for each of the three timing responses (arrival, window opening, and mean egg laying) with latitude as a smoothed predictor. We formally tested for a latitudinal reduction in pre-laying time on the breeding grounds using a GAMM in which the response was the yearly difference between arrival and mean egg laying date with latitude as a smoothed predictor.

### Overall change in breeding timing

We modeled the overall change in breeding timing over the entire time series using the mean population-year egg laying dates. We fit a single model using these mean dates as the response variable and year, latitude, and a year by latitude interaction as predictor variables. To account for spatial autocorrelation, this model was fit as a GAMM with a smoothed interaction of latitude and longitude, and it included a random slope and intercept for population identity. We visualized results of the model by extracting predictions for the overall interaction between year and latitude along with the random estimates and CI for each population.

### Breeding abundance data

Data from our monitored populations were not collected in a way that allowed us to directly assess changes in breeding abundance. To link our phenological records with broad scale changes in breeding abundance over a similar period, we used reported estimates from the Breeding Bird Survey analyzed by the United States Geological Survey. This data product includes overall linear trends for yearly percent changes in breeding population abundance for each species in the dataset. Trend estimates are derived from a hierarchical Bayesian model with cross-validation used to select the most robust model structure for each species given the available data^32^. For our purposes, we extracted the simple linear trend in tree swallow breeding season abundance between 1966-2022 for each state or province in which we had monitored populations and then joined the trend to population data. This trend is reported as a percent change in abundance per year.

Because our populations do not cover every part of the tree swallow range, the purpose of this analysis was not to reconstruct a full picture of range-wide abundance trajectories, but simply to illustrate the way that breeding abundance has changed over the last half century across the range of sites that we studied. Thus, we fit a simple linear model with the population trend as the response variable and the center latitude of each state/province for which we had monitoring data as the response variable. We note that a number of prior studies have reported similar long-term changes in tree swallow abundance across the range of locations included in our study^29,52,76^.

All analyses and figures were produced in R version 4.4.1^77^. For all smoothed terms in GAMMs we limited the number of basis dimensions to 5 to preserve biologically realistic patterns and prevent overfitting. Modeling results are reported using 95% confidence intervals throughout. Sample sizes varied for different analyses depending on what data were available from each monitored population.

## AUTHOR CONTRIBUTIONS

C. Taff, J. Shipley, D. Ardia, and M. Vitousek conceived of the project and designed the plan for data integration. C. Taff contacted contributors, coordinated data synthesis, analyzed the data, and drafted the paper. B. Tonelli, M. Tingley, and C. Youngflesh contributed modeling of arrival dates from eBird data. All authors contributed to field data collection and to writing the final manuscript.

## ACKNOWLEDGMENTS

We thank the countless technicians, students, postdocs, collaborators, and volunteers who contributed to data collection at these sites over the past 50+ years. Volunteers at the Holden Arboretum and Long Point Bird Observatory were critical to projects at those sites. We also thank the citizen scientists who contributed observations to NestWatch, eBird, and the Breeding Bird Survey along with the organizations that supported those programs.

## DATA & CODE AVAILABILITY

The cleaned and filtered dataset from contributors along with all code used for analyses and figures is publicly archived on Zenodo (https://doi.org/10.5281/zenodo.17804486). Code to process publicly available environmental data is also provided (temperature, land use, eBird, Breeding Bird Survey) and the raw data can be accessed from each project directly. NestWatch data used in the paper is publicly available from the project website (https://NestWatch.org/explore-data/NestWatch-open-dataset-downloads/).

## MATERIALS & CORRESPONDENCE

All correspondence and material requests should be addressed to Conor Taff.

## SUPPLEMENTARY FIGURES AND TABLES

**Figure S1.**
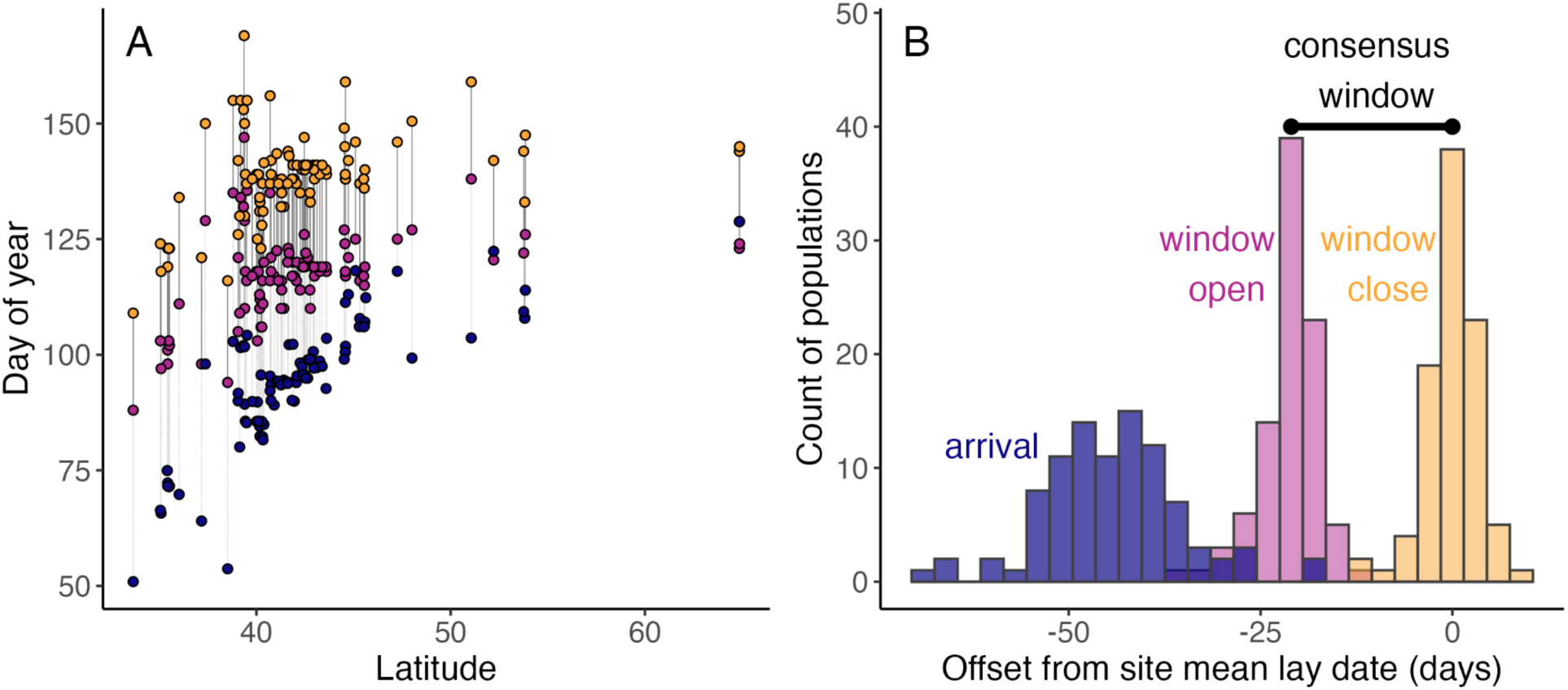
Dates of identified arrival timing and temperature sensitive periods from sliding window analyses. (A) Points show estimated arrival dates (blue), sensitive window opening date (pink), and window closing date (yellow) plotted against latitude with one point of each type for each population. Gray lines connect the arrival to window opening date and black lines connect the window opening to closing date for each population. (B) Distribution of arrival, window opening, and window closing date relative to the population-specific overall mean laying date. The identified consensus window across all populations is shown above the distributions.

**Figure S2.**
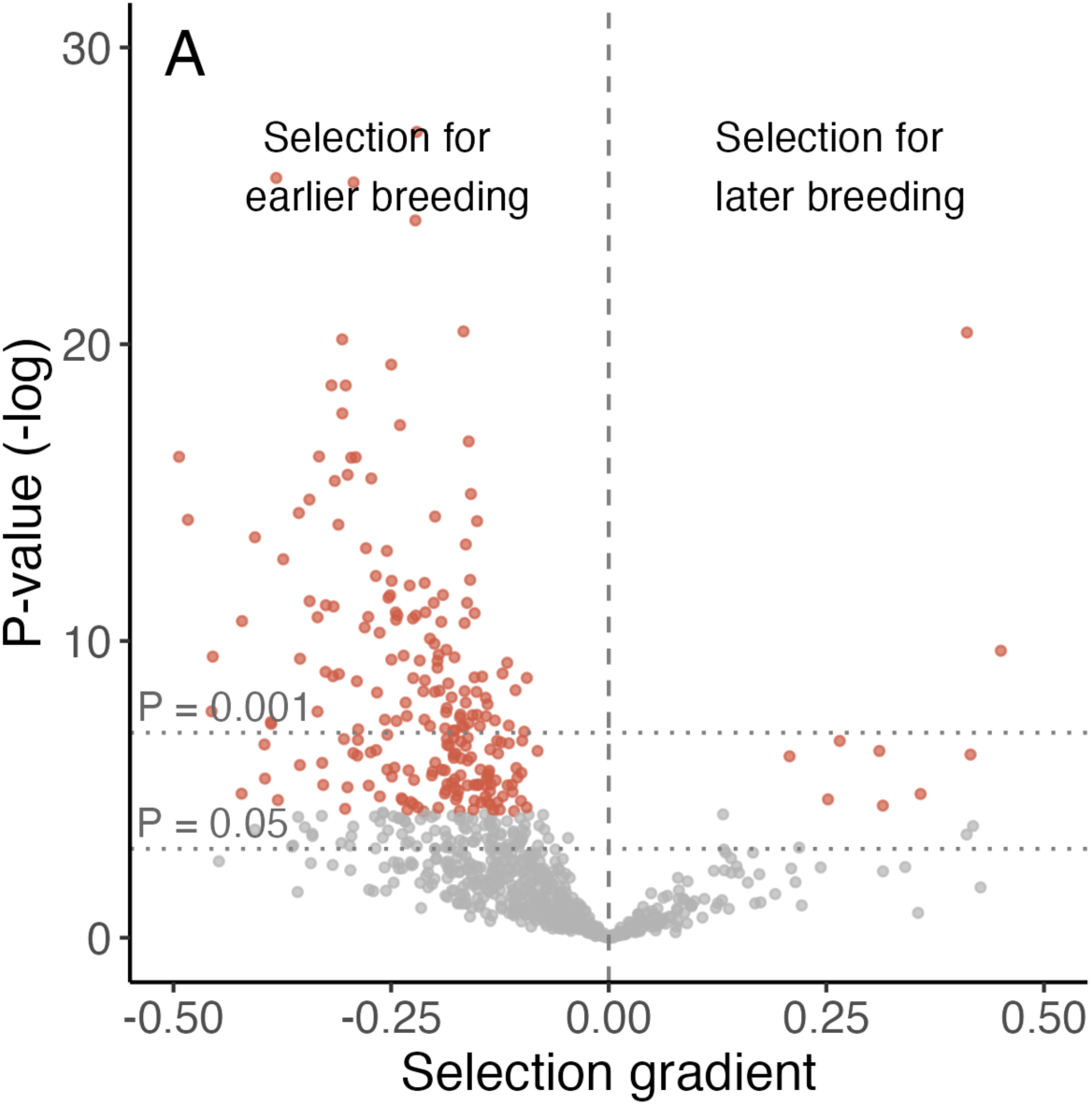
Standardized selection gradients for each population-year that met the filtering criteria. Red points were significant after the FDR correction. After correcting for multiple comparisons using the Benjamini–Hochberg procedure to control the false discovery rate, 227 population-years (27%) showed significant selection for earlier breeding, while only 16 (2%) showed significant selection for later breeding. Fourteen points with outlier values are outside of the plotting region, but were included in analyses.

**Figure S3.**
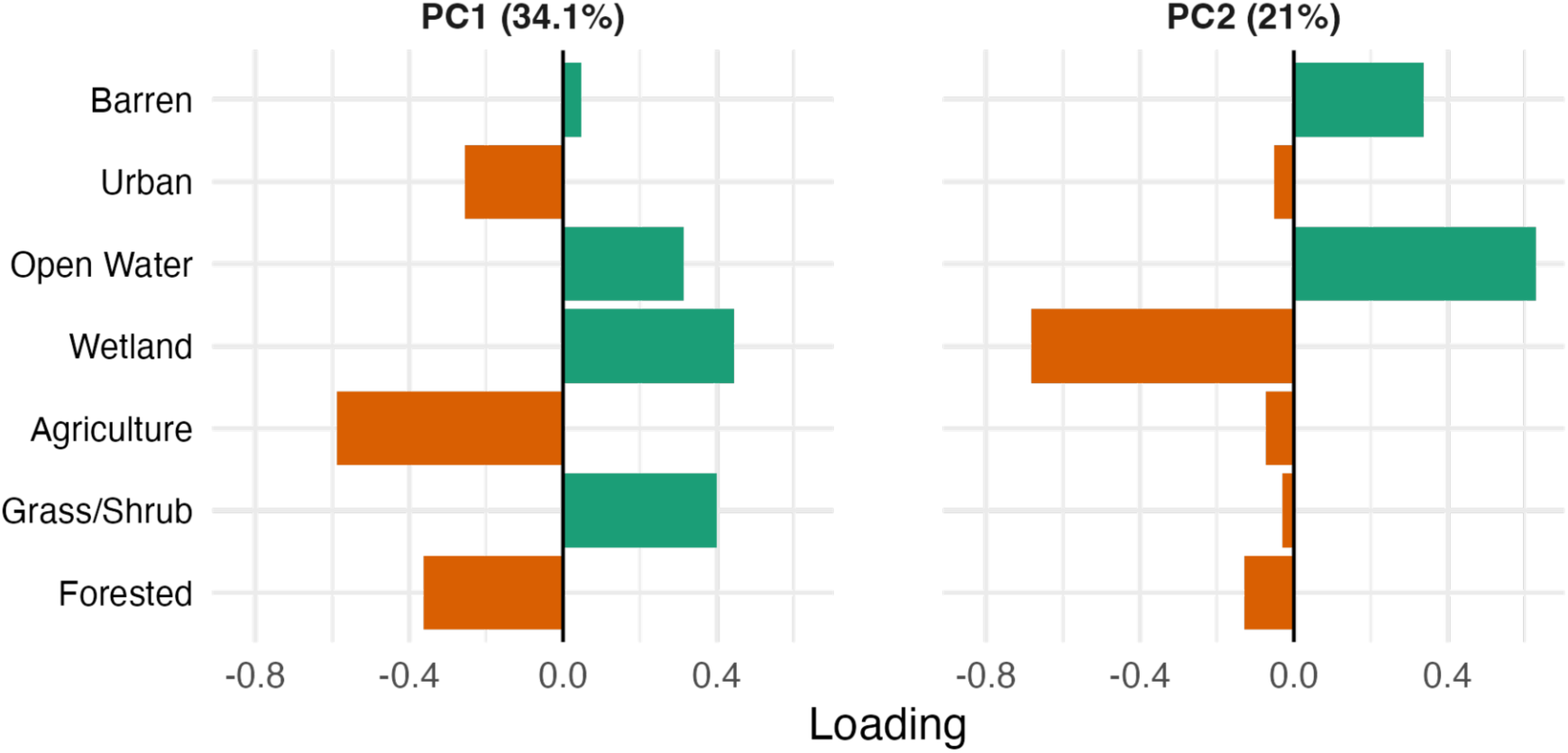
Loadings of seven land cover categories on principal component axes 1 and 2. Principal components for proportional land cover data were calculated after center log ratio transformation.

**Figure S4.**
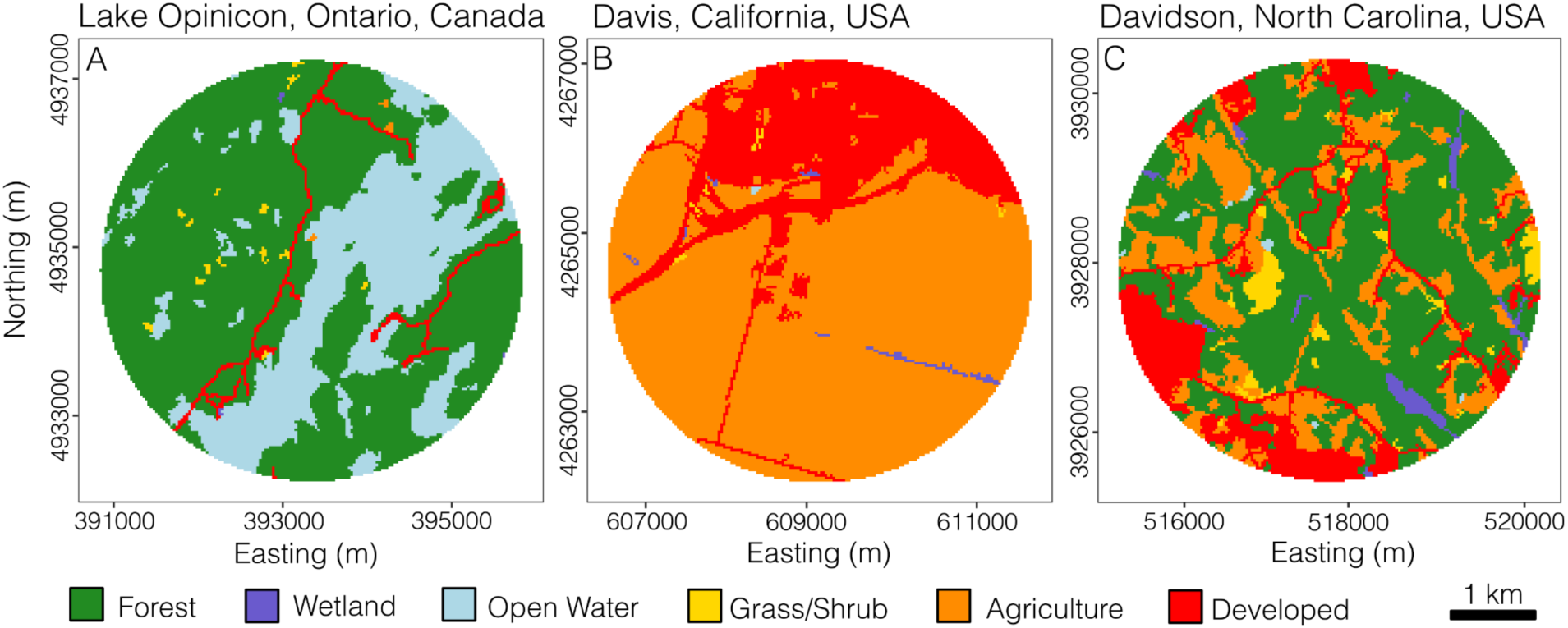
Examples of land cover in 5km diameter buffers around sites illustrating areas with high levels of forest and open water (A), agriculture and urbanization (B), and a mixture of various land cover (C). These specific populations are illustrative of the variety of land cover types that surround study sites where nest monitoring took place.

**Table S1.**
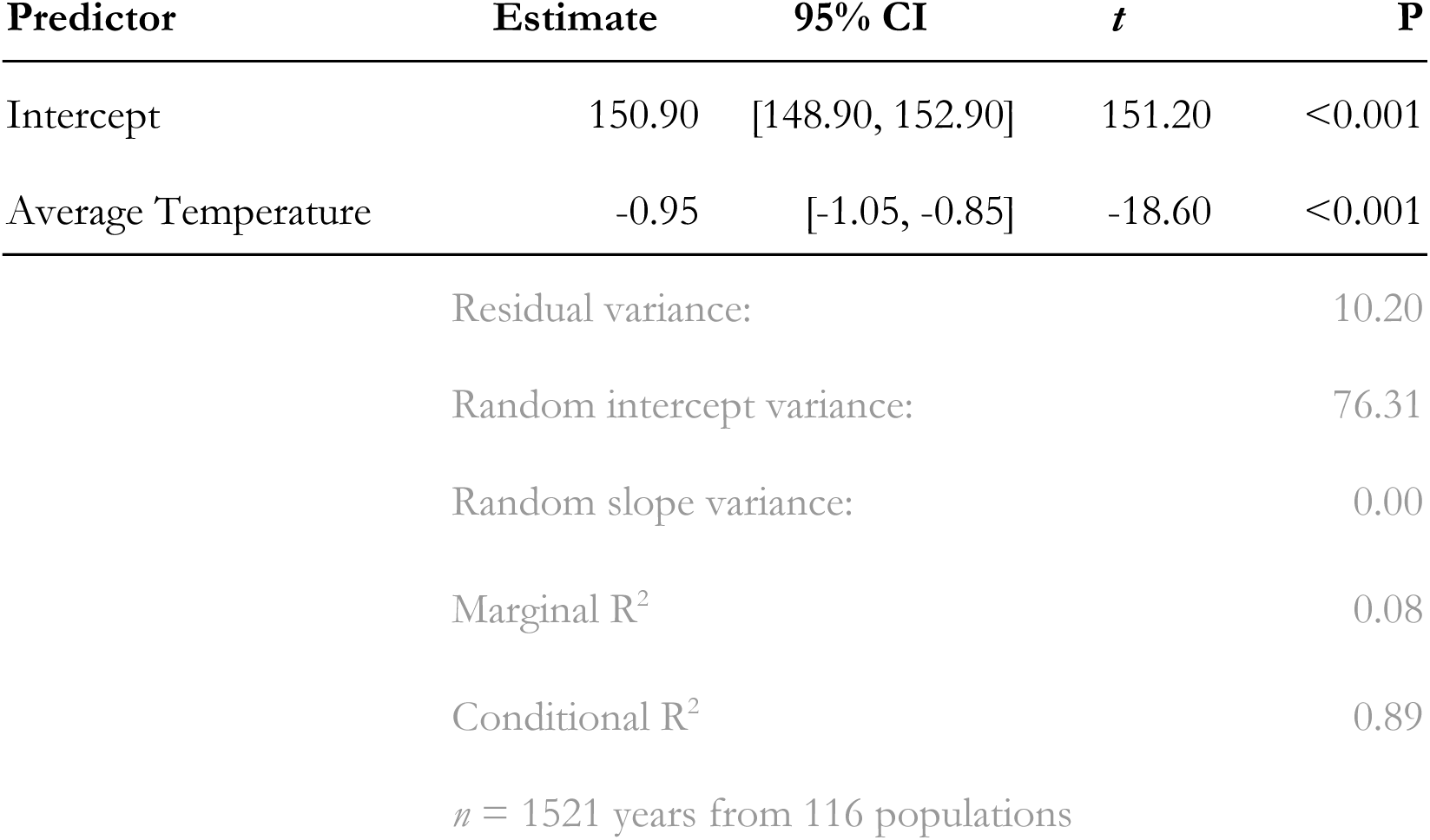
Linear mixed model results with mean population-year laying date as the response variable and random intercept and slope for population identity. Estimates are shown from the reduced model that drops the random slope.

**Table S2.** Model estimated relationships between latitude, land use, and the timing or slope of sensitive windows. Predictors were added separately as smoothed terms in GAMMs accounting for spatial autocorrelation.

| **Predictor** | **edf** | **F** | **P** |
| --- | --- | --- | --- |
| *Day of sensitive window opening (n = 116)* |  |  |  |
| latitude | 1.00 | 1.23 | 0.27 |
| land use PC1 | 1.00 | 1.13 | 0.29 |
| land use PC2 | 1.47 | 0.33 | 0.63 |
| *Day of sensitive window closing (n = 116)* |  |  |  |
| latitude | 1.00 | 0.52 | 0.47 |
| land use PC1 | 1.00 | 0.65 | 0.42 |
| land use PC2 | 1.00 | 0.71 | 0.40 |
| *Slope of response (phenological sensitivity; n = 116)* |  |  |  |
| latitude | 1.89 | 0.79 | 0.44 |
| land use PC1 | 1.00 | 0.20 | 0.66 |
| land use PC2 | 1.00 | 0.01 | 0.92 |

**Table S3.** Results of a GAMM using standardized linear selection gradient for each population-year as the response variable.

A) Linear predictors
| **Predictor** | **estimate** | **std. error** | ***t*** | **P** |
| --- | --- | --- | --- | --- |
| latitude | -0.007 | 0.003 | -2.02 | 0.04 |
| temperature anomaly | -0.002 | 0.0002 | -14.03 | <0.001 |

B) Smoothed predictors
| **Predictor** | **edf** | **F** | **P** |
| --- | --- | --- | --- |
| s(population, random) | 19.40 | 0.43 | <0.001 |
| s(longitude, latitude) | 7.69 | 2.20 | 0.02 |

$n = 826$ years from 85 populations
Deviance explained = 11.4%

**Table S4.** Results of a GAMM using standardized within-population reproductive success as the response variable.

| **Predictor** | **edf** | **F** | **P** |
| --- | --- | --- | --- |
| s(within population lay date z-score) | 3.95 | 338.01 | <0.001 |
| s(population, random) | 0.00 | 0.00 | 1.00 |
| s(longitude, latitude) | 3.00 | 1.04 | 0.37 |

$n = 63,616$ nests from 85 populations
Deviance explained = 2.1%

**Table S5.** Results of GAMMs for arrival date, sensitive window opening date, and mean egg laying date in relation to latitude and controlling for site elevation.

| **Response** | **Predictor** | **edf** | **F** | **P** |
| --- | --- | --- | --- | --- |
| *Arrival date (n = 1,212 population-years; deviance explained = 91.2%)* |  |  |  |  |
|  | s(latitude) | 2.34 | 113.02 | <0.001 |
|  | s(elevation) | 1.00 | 30.72 | <0.001 |
|  | s(population, random) | 107.40 | 23.29 | <0.001 |
| *Open window date (n = 106 population estimates; deviance explained = 76.4%)* |  |  |  |  |
|  | s(latitude) | 3.43 | 37.07 | <0.001 |
|  | s(elevation) | 3.85 | 46.91 | <0.001 |
| *Mean egg laying date (n = 1,555 population-years; deviance explained = 88.2%)* |  |  |  |  |
|  | s(latitude) | 4.00 | 93.43 | <0.001 |
|  | s(elevation) | 3.87 | 105.72 | <0.001 |
|  | s(population, random) | 100.73 | 8.25 | <0.001 |
| *Time from arrival to egg laying (n = 1,212 population-years; deviance explained = 75.3%)* |  |  |  |  |
|  | s(latitude) | 3.63 | 48.64 | <0.001 |
|  | s(elevation) | 1.00 | 22.28 | <0.001 |
|  | s(population, random) | 96.22 | 9.76 | <0.001 |

**Table S6.** Results of models illustrating latitudinal gradients in temperature change between 1950-74 vs. 2000-24, the mean laying date over time, and estimated changes in breeding season population abundance.

| **Response** | **Predictor** | **estimate** | **std. error** | ***t*** | **P** |
| --- | --- | --- | --- | --- | --- |
| *Temperature 1950-74 vs. 2000-24 (GAM; $n = 123$; deviance explained = 77.8%)* |  |  |  |  |  |
|  | latitude | 0.01 | 0.0003 | 35.35 | <0.001 |
| *Mean laying date (GAMM; $n = 1,555$ population-years; deviance explained = 88.5%)* |  |  |  |  |  |
|  | year | 0.34 | 0.14 | 2.40 | 0.016 |
|  | latitude | 7.05 | 0.59 | 11.99 | <0.001 |
|  | year*latitude | -0.01 | 0.003 | -2.99 | 0.003 |
|  | s(population, random) |  |  |  | <0.001 |
| *BBS state/province level population trend (LM; $n = 33$, $R^2 = 0.18$)* |  |  |  |  |  |
|  | latitude | -0.36 | 0.13 | -2.81 | 0.009 |

